# Phosphorylation in the *Plasmodium falciparum* proteome: A meta-analysis of publicly available data sets

**DOI:** 10.1101/2023.11.20.567785

**Authors:** Oscar J M Camacho, Kerry A Ramsbottom, Ananth Prakash, Zhi Sun, Yasset Perez Riverol, Emily Bowler-Barnett, Maria Martin, Jun Fan, Eric W Deutsch, Juan Antonio Vizcaíno, Andrew R Jones

## Abstract

Malaria is a deadly disease caused by Apicomplexan parasites of the *Plasmodium* genus. Several species of the *Plasmodium* genus are known to be infectious to human, of which *P. falciparum* is the most virulent. Post-translational modifications (PTMs) of proteins coordinate cell signalling and hence, regulate many biological processes in *P. falciparum* homeostasis and host infection, of which the most highly studied is phosphorylation. Phosphosites on proteins can be identified by tandem mass spectrometry (MS) performed on enriched samples (phosphoproteomics), followed by downstream computational analyses. We have performed a large-scale meta-analysis of 11 publicly available phosphoproteomics datasets, to build a comprehensive atlas of phosphosites in the *P. falciparum* proteome, using robust pipelines aimed at strict control of false identifications. We identified a total of 28,495 phosphorylated sites on *P. falciparum* proteins at 5% false localisation rate (FLR) and, of those, 18,100 at 1% FLR. We identified significant sequence motifs, likely indicative of different groups of kinases, responsible for different groups of phosphosites. Conservation analysis identified clusters of phosphoproteins that are highly conserved, and others that are evolving faster within the *Plasmodium* genus, and implicated in different pathways. We were also able to identify over 180,000 phosphosites within *Plasmodium* species beyond *falciparum*, based on orthologue mapping. We also explored the structural context of phosphosites, identifying a strong enrichment for phosphosites on fast evolving (low conservation) intrinsically disordered regions (IDRs) of proteins. In other species, IDRs have been shown to have an important role in modulating protein-protein interactions, particularly in signalling, and thus warranting further study for their roles in host-pathogen interactions. All data has made available via UniProtKB, PRIDE and PeptideAtlas, with visualisation interfaces for exploring phosphosites in the context of other data on *Plasmodium* proteins.

**Author Summary:** *Plasmodium* parasites continue to pose a significant global health threat, with a high proportion of the world at risk of malaria. It is imperative to gain new insights into cell signalling and regulation of biological processes in these parasites to develop effective treatments. This study focused on post-translational modifications (PTMs) of proteins, specifically phosphorylation. We conducted a meta-analysis of 11 publicly available phosphoproteomics datasets, identifying over 28,000 phosphorylated sites on *P. falciparum* proteins, using very rigorous statistics to avoid reporting false positives, and mapping to over 180,000 phosphorylation sites on other species of *Plasmodium*.

The analysis revealed distinct sequence motifs associated with different groups of phosphosites (and likely indicative of different upstream kinases), and differences in the downstream pathways regulated. Conservation analysis highlighted clusters of phosphoproteins evolving at different rates within the *Plasmodium* genus. Notably, phosphorylation was enriched in regions of proteins lacking distinct structural elements, known as intrinsically disordered regions (IDRs), which are poorly conserved across the genus – we speculate that they are important for modulating protein interactions. The findings provide valuable insights into the molecular mechanisms of *P. falciparum*, with potential implications for understanding host-pathogen interactions. The comprehensive dataset generated is now publicly accessible, serving as a valuable resource for the scientific community through UniProtKB, PRIDE, and PeptideAtlas.

## Introduction

Malaria remains a major global health burden with 247 million cases worldwide in 2021. In the same year, the World Health Organisation (WHO) has been estimated that 619,000 people died from the disease. Most malaria cases (95%) and deaths (96%) occurred in Sub-Saharan Africa. Malaria is caused by apicomplexan parasites of the *Plasmodium* genus. Of the approximately 156 named *Plasmodium* species, only five have been found to infect humans: *P. falciparum*, *P. vivax*, *P. ovale*, *P. malariae*, and *P. knowlesi*. *Plasmodium* is transmitted from one human to another by female *Anopheles* mosquitos with the exception of *P. knowlesi*, which is believed to be zoonotic i.e. transmission happens from macaques to humans in southeast Asia, where macaques have been previously infected by *Anopheles* mosquitos [1]. Most severe cases and deaths from malaria are caused by *P. falciparum* infections, endemic to Sub-Saharan Africa.

The *P. falciparum* life cycle requires two hosts, an *Anopheles* mosquito (around 40 species of *Anopheles* can transmit *P. falciparum* [2]) and a human host. Extracellular sporozoites are transmitted to the human dermal tissue during a blood meal. Once they reach the liver they replicate and develop into merozoites and get released into the peripheral blood. There, merozoites break into erythrocytes and replicate, causing most malaria symptoms. A small proportion of parasites will develop into gametocytes with sexual attributes, which, closing the cycle, are transmitted back to mosquitoes where sporozoites are formed from gametocytes [3].

Post-translational modifications (PTMs) of proteins are critically important as they can act as molecular switches. PTMs and specifically phosphorylation has been shown to dynamically change, for example, between extra and intraerythrocytic life-cycle stages suggesting protein sets and pathways with roles in cell invasion [4]. Across stages in the *Plasmodium* intraerythrocytic asexual cycle, a study has reported changes in abundance, defined as peak changes of at least 1.5 fold between stages, for 34% of identified proteins and 75% of phosphorylation sites [5]. Interruption and obstruction of interactions among these proteins could constitute efficient treatments against malaria.

All stages in the *Plasmodium* life cycle have the potential to generate targets for vaccines. For example, transmission-blocking vaccines preventing mosquito infection or interfering in *Plasmodium* sexual stages; pre-erythrocytic stage by disabling the ability of merozoites to reach the human liver or replicate there; or targeting interactions at the blood stage by suppressing their ability to enter erythrocytes or replicate [6]. The most advanced vaccine (RTS,S), targets the *P. falciparum* circumsporozoite surface protein (PfCSP) [7-9]. Artemisinin-based combination treatments (ACTs) have proved effective for treating *P. falciparum* malaria [10, 11] but increasing *Plasmodium* drug resistance has been observed, highlighting the importance of development of new drugs [12].

There are several useful online resources to support research in *Plasmodium*. PlasmoDB [13, 14] stands out, it is part of the Eukaryotic Pathogen, Vector and Host Informatics Resources [15] (VEuPathDB), which has been running since 2004, collating genome, functional genomic and phenotypic data sets for multiple *Plasmodium* species. PlasmoDB hosts 20 *Plasmodium falciparum* annotated genomes, including the canonical reference P. *falciparum* clone 3D7 [16]. A search of the *P. falciparum* proteome in the UniProt Knowledgebase (UniProtKB) [17], the world’s most popular protein knowledge-base, returns 18 proteomes linked to different isolates of countries of origin, which are mostly well synchronised with PlasmoDB.

Tandem mass spectrometry (MS/MS) is most often used in large scale phosphosite identification and localisation studies [18]. Protein samples are purified and enzymatically digested, typically using trypsin. Samples are then enriched for phosphorylated peptides using reagents such as TiO2, or other metal ions that promote phosphate binding. Then liquid chromatography is used to separate peptides that are subsequently fragmented and analysed by MS/MS. Results from MS analyses can then be compared against protein sequence databases, with and without the mass shift for phosphorylation, via one of the many available search algorithms [19, 20]. Algorithms provide identification of peptides and localisation of PTM sites in those peptides with scores aiming to reflect the level of confidence that those identifications are correct. Score thresholding is used to select a subset of what are expected to be the most confident findings. However, score thresholding does not provide information about the global false discovery rate (FDR) of peptides, or the global false localisation rate (FLR) of the phosphosites within those peptides. Absence of objective calculation of FLR in phophosite localisation studies hinders comparisons among studies, as it is not possible to establish a common quality threshold among results from different studies. To overcome this problem, we have recently published an approach that allows estimation of global site-level FLR, by including a decoy amino acid, specifically Alanine, for phosphorylation searches (which cannot be modified) as a search parameter to compete against targets sites (S, T or Y), i.e. the pASTY method [21]. An important benefit of pASTY searches is that it allows combination of results from multiple studies as FLR can provide objective comparable thresholds [22].

For *P. falciparum,* PlasmoDB provides information on 16,118 phosphorylation sites, although phosphorylation sites have been loaded from multiple publications over the last 10 years. In our previous work examining databases containing human phosphosites, we estimated that there is a high proportion of false positive sites recorded, due to historically inadequate statistics associated with detection of sites by MS [23] and, prior to [22], lack of methods for calculating adequate statistics for controlling the FLR across studies.

In this work, we aim to provide a high-quality mapping of *P. falciparum* phosphosites via a large-scale re-analysis of public phospho-enriched studies, underpinned by robust analysis pipelines enabling meta-analyses with FLR control within and across studies’ results. This analysis is part of the “PTMeXchange” initiative, which is re-analysing phospho-enriched data sets and depositing results into the proteomics resources PRIDE [24], PeptideAtlas [25] and UniProtKB. Results from downstream analysis in the most confident phosphosites are also reported in this manuscript, including analysis motifs centred on phosphosites and pathway enrichment analysis for these motifs. We examine phosphorylation site conservation between our reference isolate 3D7 (*P. falciparum*) and other species of the *Plasmodium* genus, as well as investigating the structure and disordered regions of phosphoproteins.

## Results

### Phosphosite identification

The counts of peptide-spectrum matches (PSMs), and PSM-sites passing the 1% FDR threshold are displayed in Table 1 as well as the number of overall PSM-sites at 1% and 5% FLR. Next, data were collapsed to peptidoform-site level *i.e.* removing redundancy caused by the common occurrence of multiple PSMs identifying the same peptidoform. A peptidoform is defined as a unique sequence of amino acids with specific modifications. For example, two identical peptide sequences but with modifications at different positions in the sequence are different peptidoforms. Table 1 also displays the number of peptidoform-sites and protein-sites (accepting the mapping from a peptide sequence to all proteins it can be found in, assuming tryptic cleavage) at 1% and 5% FLR, with these counts separated for *P. falciparum* and human matches. Note that FLR threshold counts only considered PSMs that passed 1% FDR. The data is unequally distributed between the 11 studies, with four studies (PXD012143, PXD015833, PXD020381, PXD026474) contributing significantly more to the overall number of sites at every stage of the analysis. In fact, nearly 70% of all *P. falciparum* protein-sites at 5% FLR come from these four studies.

**Table 1.** From left to right: PSM count at 1% FDR, PSM-site count at 1% FDR, overall PSM-site count at 1% and 5% FLR, P. falciparum peptidoform-site count at 1% and 5% FLR, human peptidoform-site count at 1% and 5% FLR, P. falciparum protein-site count at 1% and 5% FLR, human protein-site count at 1% and 5% FLR.

|  |  |  | Overall |  | Plasmodium |  | Human |  | Plasmodium |  | Human |  |
| --- | --- | --- | --- | --- | --- | --- | --- | --- | --- | --- | --- | --- |
|  | 1% FDR PSM | 1% FDR PSM-site | 1% FLR PSM-Site | 5% FLR PSM-Site | 1% FLR Peptidoform-site | 5% FLR Peptidoform-site | 1% FLR Peptidoform-site | 5% FLR Peptidoform-site | 1% FLR Protein-site | 5% FLR Protein-site | 1% FLR Protein-site | 5% FLR Protein-site |
| PXD000070 | 42054 | 46064 | 6206 | 12375 | 564 | 1265 | 44 | 103 | 398 | 886 | 27 | 69 |
| PXD001684 | 14152 | 14179 | 5648 | 12015 | 901 | 1291 | 48 | 56 | 785 | 1110 | 45 | 63 |
| PXD002266 | 38618 | 45004 | 10342 | 18609 | 1609 | 3298 | 68 | 150 | 1147 | 2421 | 47 | 118 |
| PXD005207 | 21967 | 24045 | 4584 | 8105 | 2367 | 3386 | 188 | 296 | 1364 | 1884 | 105 | 174 |
| PXD009157 | 22449 | 24653 | 4134 | 7049 | 2179 | 3353 | 83 | 151 | 1291 | 2023 | 31 | 69 |
| PXD009465 | 24435 | 37096 | 7812 | 10694 | 3905 | 5382 | 101 | 159 | 2513 | 3534 | 70 | 122 |
| PXD012143 | 270306 | 340996 | 93438 | 144256 | 26893 | 39487 | 633 | 1139 | 10597 | 15797 | 350 | 662 |
| PXD015093 | 28693 | 30469 | 5597 | 12187 | 1156 | 2760 | 123 | 274 | 583 | 1242 | 64 | 149 |
| PXD015833 | 348842 | 392870 | 78345 | 134576 | 22288 | 43134 | 1291 | 3005 | 6439 | 11897 | 442 | 1060 |
| PXD020381 | 228888 | 278203 | 67457 | 105938 | 17604 | 31622 | 361 | 832 | 7331 | 13050 | 173 | 483 |
| PXD026474 | 154644 | 168133 | 57786 | 100714 | 8772 | 14611 | 3334 | 5640 | 4657 | 7426 | 1549 | 2657 |

Figure 1A shows the Gold-Silver-Bronze (GSB, see Methods) quality categorisation for *P. falciparum* protein-sites matching to *P. falciparum* (left panel) and human (right panel) proteins. If a sequence, and therefore the sites within the sequence, match to more than one protein sequence, these sites were counted for all mapping proteins in the proteome. In total, there were 28,495 protein-sites in *P. falciparum* classified as GSB (5% FLR) of which 9,587 were Gold (seen in 2 or more studies at <1% FLR), 8,513 Silver (1 study at <1% FLR), and 10,395 Bronze (>=1% FLR and <5% FLR). While for Human (Figure 1A right), there were 4,239 protein-sites identified, with 446 Gold, 1,788 Silver and, 2,005 classified as Bronze. 25,758 and 3,529 sites were unique (matching to one protein only), and 351 and 284 non-unique sites for *P. falciparum* and Human respectively, considering only sites at 5% FLR (Figure 1B). The number of GSB unique protein-sites were 26,028 in *P. falciparum* and 3,732 in Human (Figure 1C), with 81 protein-sites matching to both species (this calculation only considers a protein match per sequence and site). Only 1.35% of all protein-sites in our *Plasmodium* analysis mapped to more than 1 protein with less than 0.05% mapping to more than two proteins. Interestingly, there was a peptidoform mapping the same site evidence to 36 proteins from the PfEMP1 family, due to very high sequence similarity amongst this gene family (Figure 1D). All Gold, Silver and Bronze phosphosites can be found in Supp File 1.

**Figure 1.**
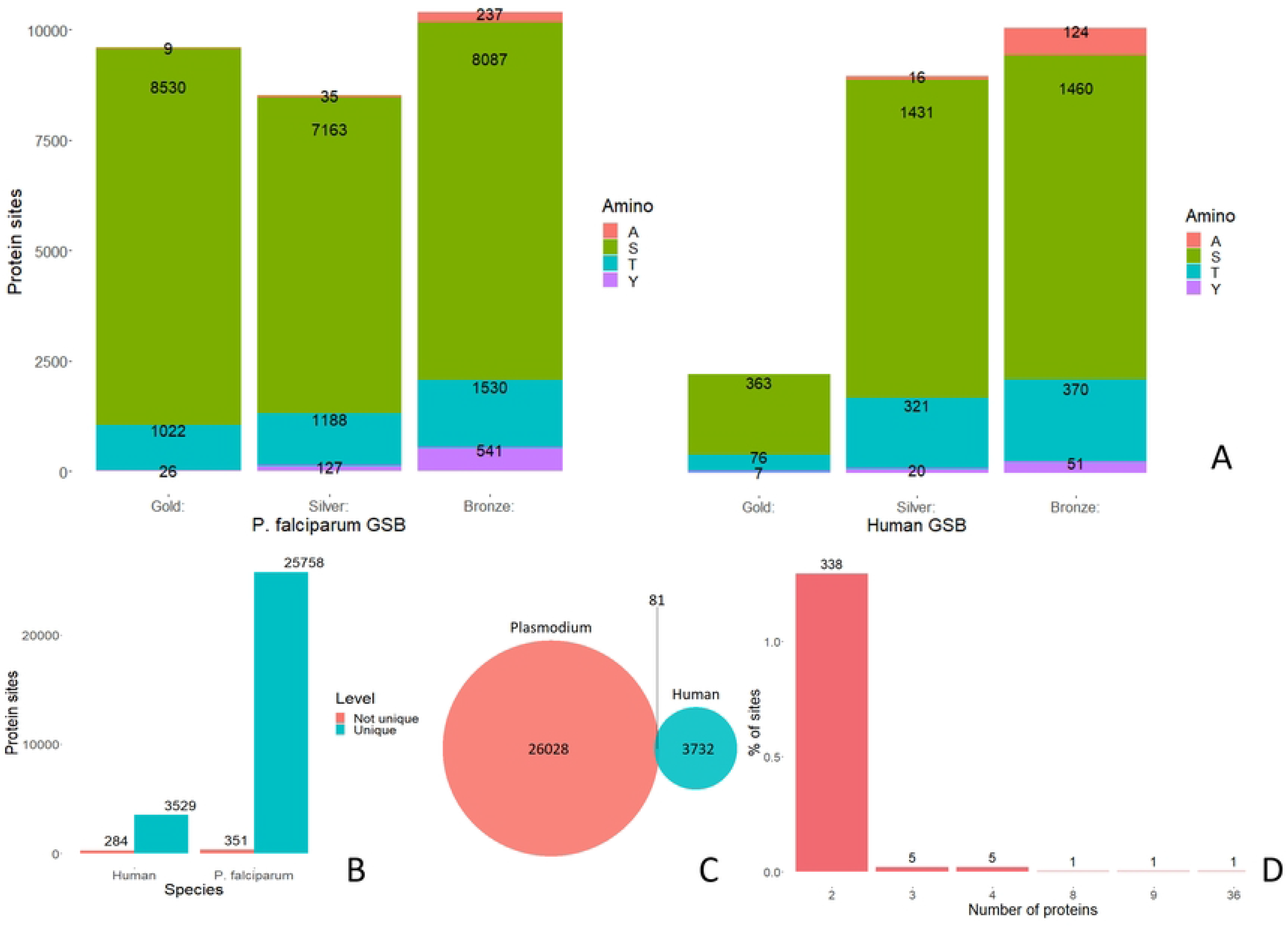
A: The count of protein-sites classified as Gold-Silver-Bronze for *P. falciparum* (left) and Human (right), sites coloured by phosphorylated amino acid. This includes all potential locations for the identified sites when peptides match to more than one location or protein and decoy matches to Alanine. B: Number of unique or not unique protein-sites identified within each species among sites at 5% FLR, where not unique are those peptidoforms that map to more than one protein. C: Number of protein-sites, where sites are mapped to a single protein, for each species and common to both. D: Proportion of sites matching to more than one protein.

### Motif and pathway enrichment analysis

We investigated if there were overrepresented sequence motifs centred on 5% FLR phosphosites. Motif analysis was performed for *P. falciparum* by comparing 15mer peptides centred on S, T and Y phosphosites against a background of 15mer peptides centred on all STY sites, phosphorylated or not. The analysis returned 107 statistically significant motifs, of which 65, 30 and 12 were centred on S, T and Y, respectively (Supp Figure 1 and Supp File 1).

The most common significant motifs (group 1 on Figure 2) were those with S and T combining with E and D in different positions such: [ST]D, D[ST], [ST].[DE], [ST]…D, and [ST]N[DE], N[ST].[DE] or N.[ST] [DE]. There were also other common motifs to S and T not containing D or E like N[ST], K..[ST], R..[ST]. Among other potentially interesting features of these motifs, only K was found in motifs at positions - 7, -6, +6 and +7, with significant motifs for K…..DS, K….DS and SD….K, SD…..K, SN…..K. Motifs with K relatively distant to the target site may be due to a preference for particular kinases or an artefact related to tryptic cleavage enriching for lysines to be present somewhere in many detected peptides. There were only six 3-amino acid motifs with a P in the +1 position, which can be very common for other species. Many of these motifs agreed with those found by Peace *et al.* [5], which can be expected as their MS data was included in this analysis. Treeck *et al.* [26] also found many of these motifs in *P. falciparum* (S[DE].E,SD., K..S.D, [KR]..S, S[DE], SN, DS).

**Figure 2.**
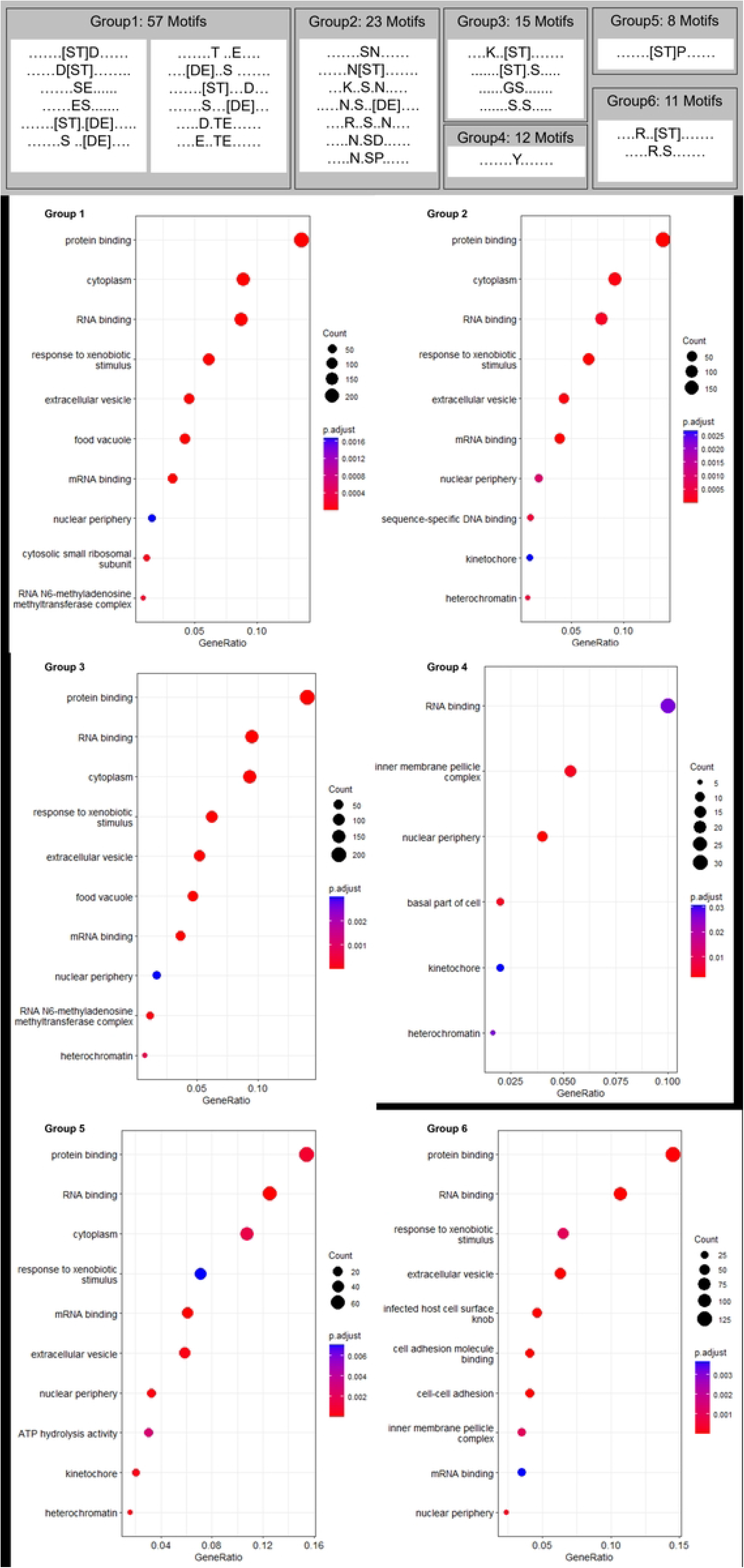
Statistically significant GO terms for motifs surrounding phosphorylated sites, grouped by similar motifs, formed by similar groups of amino acids.

Motifs were independently analysed to investigate functional enrichment of the proteins in which specific motifs were found. Summing across all motif results, 39 different GO terms were found to be statistically significant. Table 3 contains significant GO terms for motifs at least 4-fold enriched versus the background, and the full analysis for all motifs can be found in Supp File 2. It is worth noting that the RSF.D motif gave highly significant matches to several GO terms. However, this result is an unusual artefact of the protocol. *P. falciparum* has a 65 gene family, in which all members are called “PfEMP1”, with highly related protein sequences. Phosphosites are mapped to all positions where a peptidoform can be found, typically resulting in the vast majority of sites mapping to a single protein (counts for the few exceptions are shown in Figure 1D). Phosphosites identified in PfEMP1 mapped to 36 different proteins, which gives an apparently extremely significant signal under motif-GO analysis, since the PfEMP1 proteins all contain the same motif and are also all mapped to the same GO terms.

Under the hypothesis that similar motifs may indicate proteins being involved in similar processes, we identified six groups based on the amino acids forming those motifs (Figure 2). Our analysis showed that besides general agreement on high-level functional terms such as “protein binding”, “cytoplasm” or “RNA binding” there were also group specific GO terms pointing to some functional specificity among motifs’ groups. There was almost complete agreement in significant terms between groups 1 to 3 which could suggest there is overlap in the signalling cascades with regards to downstream effects, or the method may not be sensitive enough to draw more specific terms allowing differentiation. Groups 4 to 6 returned more diverse GO terms, with group 6 likely conditioned by the PfEMP1 protein family as previously explained.

As an alternative and more objective approach for grouping motifs, we took a subset of motifs with a 4-fold change over background, *i.e.* the most strongly over-represented motifs, and performed an enrichment analysis for all *P. falciparum* proteins where those phosphorylation motifs can be found. Figure 3A shows a heatmap of the p-values for the GO terms associated with proteins containing these phosphorylation motifs. These motifs did not seem to form strong clusters according to these terms, although some small groups (similar pairs clustering together) could be observed. In a few cases, there was some overlap in protein sets carrying motifs, for example SD.E and SD.E..K – the latter being matched to a subset of the proteins matched by the former. However, the motifs K..T.E and K..T.D are mutually exclusive, yet proteins carrying these phosphosite motifs can be seen to be acting in many of the same signalling pathways. Other examples of “pairing” on the dendrogram are motifs N.SP and N..SP, and E..T.E and E…TE. It is possible that these pairs of phospho-motifs are recognised by the same kinase, or that there are closely related kinases within the same family that are involved within the same types of downstream pathways. Another “pair” of phospho-motifs is K..TP and TDD – this latter example is surprising, since it would typically be assumed that S/TP and S/TD phosphosites are regulated by different kinase families. It is possible that there is crosstalk between two kinases, although the evidence here is not sufficient to form any strong conclusions.

**Figure 3.**
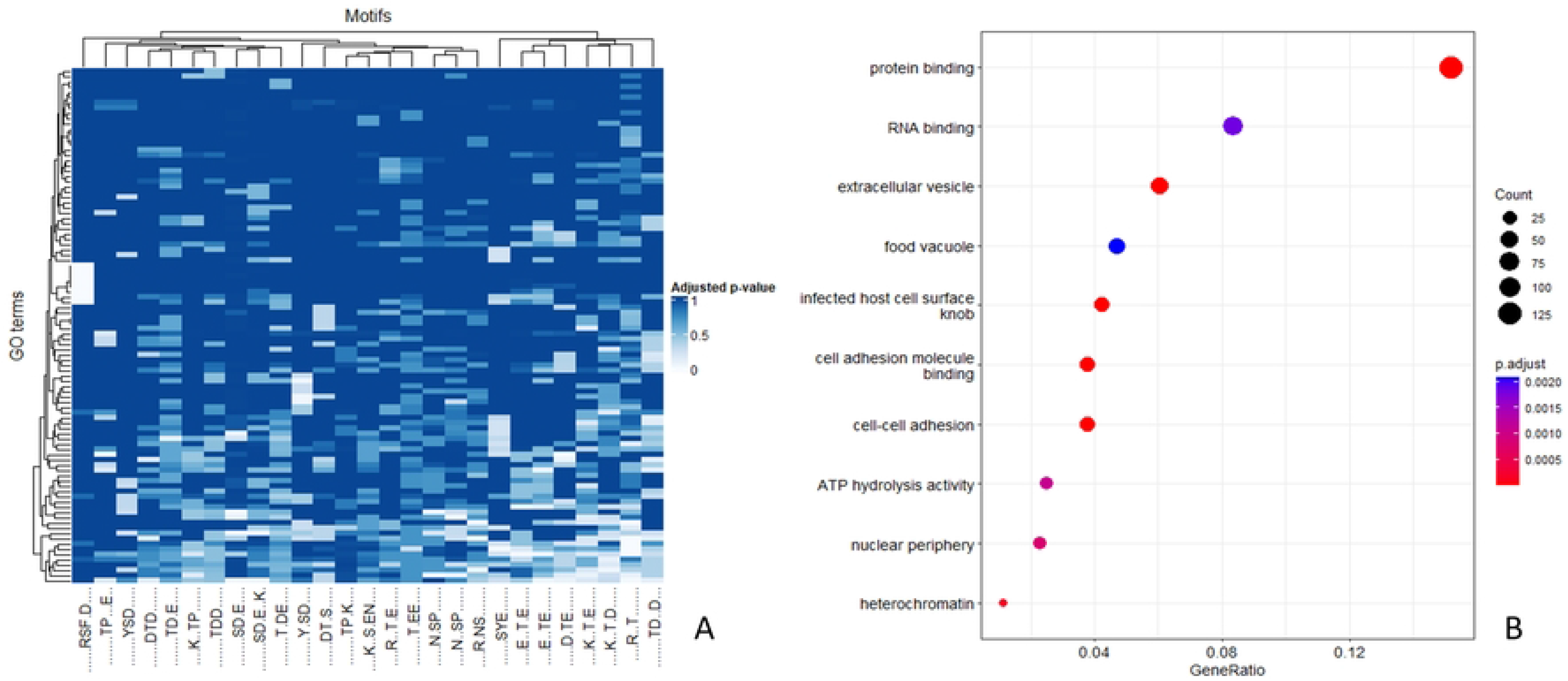
A: Heatmap of the adjusted p-values resulting from an enrichment analysis of the proteins where phosphorylation motifs (on the x-axis) were found (only motifs with fold change over background above 4 are considered). B: Significant GO Terms (p-adjusted <=0.05) for the genes in which motifs with fold change over background above 4 could be found.

Therefore, in general, the heatmap suggests some differences in functionality between proteins containing different phosphorylation motifs. However, when analysed as a group, 10 GO terms summarised the functional processes for the subset of genes where these phosphorylated motifs were found (Figure 3B).

The *P. falciparum* proteome has around 90 protein kinases – 89 are annotated in PlasmoDB with the InterPro protein kinase domain term. An analysis by Adderley *et al.* [27] identified 98 protein kinases in isolate 3D7, by including keyword searching in addition to InterPro domain searching. From their list of kinases, there several sequences annotated as pseudokinases or having kinase-like domains (PF3D7_0424700, PF3D7_0708300, PF3D7_0724000, PF3D7_0823000, PF3D7_1106800, PF3D7_1321100, PF3D7_1428500) and two pseudogenes (PF3D7_0731400 and PF3D7_1476400). Adderley *et al.* [27] classified these kinases into families, using an HMM (Hidden Markov Models)- based technique. Supp Table 2 shows the counts of 3D7 kinases classified into families, with additional notes on potential kinase motifs for these families based upon information on kinase motifs found in humans from [28], with the caveat that we cannot be sure that even if orthologous kinases exist between humans and *P. falciparum* that they recognise the same motifs. Accurate prediction of kinase-substrate relationships is not straightforward without high-quality experimental data, which is lacking in *P. falciparum.* FIKK kinases are unique to Apicomplexa [29], and seem to have an important role in host invasion (such as phosphorylation of erythrocyte proteins), but otherwise little is known about the motifs for their targets. The publication associated with dataset PXD015833 specifically investigated substrate specificity and concluded that some of the FIKK family have a preference for a basic motif near to pS/pT site – with arginine enriched in the minus position (group 6 in our analysis) [30]. We could speculate that the motifs groups identified in Figure 2 are driven by different kinase groups in Supp. Table 2, but without new experimental data, robust conclusions are not possible. This is an area requiring further work.

**Table 2.** Summary results and sample used for each study included in this analysis.

| Study ID | Summary | Sample | Ref |
| --- | --- | --- | --- |
| PXD000070 | "We analysed the Plasmodium falciparum schizont phosphoproteome using for the first time, a data-dependent neutral loss-triggered-ETD (DDNL) strategy and a conventional decision-tree method. ... combination of Mascot Percolator and turbo-SLoMo represents a robust workflow for data analysis using CID and ETD fragmentation." | *P. falciparum* 3D7 was cultured in 2.5–5% O+ human erythrocytes. Infected erythrocytes were processed for analysis. | [39] |
| PXD001684 | "phosphoproteome analysis of extracellular merozoites revealing 1765 unique phosphorylation sites including 785 sites not previously detected in schizonts." | *P. falciparum* 3D7 merozoites cultured *in vitro* | [4] |
| PXD002266 | "employing chemical and genetic tools in combination with quantitative global phosphoproteomics, we identify the phosphorylation sites on 69 proteins that are direct or indirect cellular targets for PfPKG. These PfPKG targets include proteins involved in cell signalling, proteolysis, gene regulation, protein export and ion and protein transport, indicating that cGMP/PfPKG acts as a signalling hub that plays a central role in a number of core parasite processes." | *P. falciparum* blood stage 3D7 (wild type)-, PKGT618Q- and CDPK1-HA- parasites were cultured *in vitro*, developing trophozoite and schizont stage parasites. For the time-course experiments, parasites were synchronized by two rounds of sorbitol treatment—first treatment when the parasites culture was at late ring/trophozoite stage and second when the parasite culture contained schizonts and ring stage parasites. | [40] |
| PXD005207 | "PfCDPK1 is critical for asexual development of Plasmodium falciparum, ... this kinase is critical for the invasion of host erythrocytes. Furthermore, using a multidisciplinary approach involving comparative phosphoproteomics we gain insights into the underlying molecular mechanisms." | *P. falciparum* 3D7 asexual blood stages were cultured in O+ erythrocytes. *P. falciparum* CDPK1- 3HA-DD parasites were cultured in the presence of 2.5 nM WR99210 and 0.25 μM Shld-1. | [41] |
| PXD009157 | "Plasmodium falciparum phosphodiesterase $\beta$ (PDEβ) hydrolyses both cAMP and cGMP and is essential for blood stage viability. Conditional gene disruption causes a profound reduction in invasion of erythrocytes and rapid death of those merozoites that invade." | *P. falciparum* erythrocytic stages were cultured in human A+ erythrocytes | [42] |
| PXD009465 | "To better understand PfPK7-regulated phosphorylation events, we performed isobaric tag-based quantitative comparative phosphoproteomics of the schizont and segmenter stages from wild-type and pfpk7- parasite lines." | Both the *P. falciparum* parasite pfpk7- line and its parental 3D7 clone were grown in RPMI 1640 culture medium supplemented with A+ erythrocytes and 0.5% Albumax | [43] |
| PXD012143 | "investigated the role of cAMP in asexual blood stage development of Plasmodium falciparum through conditional disruption of adenylyl cyclase beta (ACβ) and its downstream effector, cAMP-dependent protein kinase (PKA)." | Both the *P. falciparum* parasite pfpk7- line and its parental 3D7 clone were grown in RPMI 1640 culture medium supplemented with A+ erythrocytes and 0.5% Albumax | [44] |
| PXD015093 | "global phosphoproteomic analysis of merozoites to identify signaling pathways that are activated during invasion." | *P. falciparum* 3D7 blood stages were cultured *in vitro* to generate merozoites | [45] |
| PXD015833 | Reports "species-specific phosphorylation of erythrocyte proteins by *P. falciparum*, but not by Plasmodium knowlesi, which does not export FIKK kinases." | *P. falciparum* parasites were cultured in complete media and *P. knowlesi* parasites were cultured in CM supplemented with 10% human serum | [30] |
| PXD020381 | "identify a multipass membrane protein, ICM1, with homology to transporters and calcium channels that is tightly associated with PKG in both asexual blood stages and transmission stages. Phosphoproteomic analyses reveal multiple ICM1 phosphorylation events dependent on PKG activity." | *P. falciparum* lines were maintained in human RBCs in RPMI 1640 containing AlbuMAX II | [46] |
| PXD026474 | "understand - CDPKs - role in human parasite transmission from the host to the mosquito vector and thus investigated the role of the human-infective parasite Plasmodium falciparum CDPK4 in the parasite life cycle. *P. falciparum* cdpk42 parasites created by targeted gene deletion showed no effect in blood stage development or gametocyte development. However, cdpk42 parasites showed a severe defect in male gametogenesis and the emergence of flagellated male gametes." | *P. falciparum* NF54 and Pfcdpk42 parasites were cultured as asexual blood stages according to standard procedures and received complete RPMI medium. Gametocytes were also cultured | [47] |

**Table 3.** Statistically significant GO terms for the subset of motifs surrounding phosphorylated sites. Analyses were carried out independently for each motif and includes only motifs with at least 4 fold change over background.

| ID | Description | GeneRatio | BgRatio | pvalue | p.adjust | qvalue | Count | motif |
| --- | --- | --- | --- | --- | --- | --- | --- | --- |
| GO:0034399 | nuclear periphery | 4/52 | 35/3454 | 0.0016 | 0.0374 | 0.0367 | 4 | .....SD.E..K. |
| GO:0005515 | protein binding | 13/52 | 354/3454 | 0.0017 | 0.0374 | 0.0367 | 13 | .....SD.E..K. |
| GO:0045178 | basal part of cell | 4/144 | 11/3454 | 0.0007 | 0.0314 | 0.0302 | 4 | ....R.NS..... |
| GO:0048870 | cell motility | 4/144 | 12/3454 | 0.0011 | 0.0314 | 0.0302 | 4 | ....R.NS..... |
| GO:0009410 | response to xenobiotic stimulus | 14/144 | 145/3454 | 0.0024 | 0.0470 | 0.0451 | 14 | ....R.NS..... |
| GO:0000792 | heterochromatin | 5/233 | 11/3454 | 0.0004 | 0.0209 | 0.0197 | 5 | .....SD.E.... |
| GO:0005515 | protein binding | 40/233 | 354/3454 | 0.0005 | 0.0209 | 0.0197 | 40 | .....SD.E.... |
| GO:0034399 | nuclear periphery | 8/233 | 35/3454 | 0.0018 | 0.0472 | 0.0446 | 8 | .....SD.E.... |
| GO:0050839 | cell adhesion molecule binding | 30/36 | 54/3454 | 5.59E-53 | 6.71E-52 | 2.35E-52 | 30 | .....RSF.D.... |
| GO:0098609 | cell-cell adhesion | 30/36 | 57/3454 | 5.57E-52 | 3.34E-51 | 1.17E-51 | 30 | .....RSF.D.... |
| GO:0020030 | infected host cell surface knob | 30/36 | 61/3454 | 9.17E-51 | 3.67E-50 | 1.29E-50 | 30 | .....RSF.D.... |
| GO:0020002 | host cell plasma membrane | 30/36 | 214/3454 | 1.10E-31 | 3.30E-31 | 1.16E-31 | 30 | .....RSF.D.... |
| GO:0020013 | modulation by symbiont of host erythrocyte aggregation | 29/36 | 191/3454 | 2.49E-31 | 5.98E-31 | 2.10E-31 | 29 | .....RSF.D.... |
| GO:0020035 | adhesion of symbiont to microvasculature | 28/36 | 172/3454 | 7.88E-31 | 1.58E-30 | 5.53E-31 | 28 | .....RSF.D.... |
| GO:0020033 | antigenic variation | 29/36 | 202/3454 | 1.40E-30 | 2.41E-30 | 8.44E-31 | 29 | .....RSF.D.... |
| GO:0046789 | host cell surface receptor binding | 7/36 | 37/3454 | 5.95E-08 | 8.93E-08 | 3.13E-08 | 7 | .....RSF.D.... |
| GO:0043565 | sequence-specific DNA binding | 4/77 | 17/3454 | 0.0004 | 0.0178 | 0.0160 | 4 | ....N..SP..... |
| GO:0009410 | response to xenobiotic stimulus | 10/75 | 145/3454 | 0.0009 | 0.0484 | 0.0416 | 10 | ....K..T.D.... |
| GO:1903561 | extracellular vesicle | 7/57 | 91/3454 | 0.0006 | 0.0221 | 0.0206 | 7 | ....E..T.E.... |
| GO:0042393 | histone binding | 2/19 | 12/3454 | 0.0018 | 0.0274 | 0.0212 | 2 | .....TP...E.. |
| GO:1903561 | extracellular vesicle | 20/209 | 91/3454 | 2.51E-07 | 1.88E-05 | 1.72E-05 | 20 | ....R..T..... |
| GO:0003723 | RNA binding | 27/209 | 199/3454 | 4.23E-05 | 0.0014 | 0.0013 | 27 | ....R..T..... |
| GO:0020020 | food vacuole | 17/209 | 98/3454 | 5.83E-05 | 0.0014 | 0.0013 | 17 | ....R..T..... |
| GO:0045178 | basal part of cell | 5/209 | 11/3454 | 0.0002 | 0.0049 | 0.0045 | 5 | ....R..T..... |
| GO:0048870 | cell motility | 5/209 | 12/3454 | 0.0004 | 0.0064 | 0.0059 | 5 | ....R..T..... |
| GO:0005515 | protein binding | 35/209 | 354/3454 | 0.0018 | 0.0236 | 0.0216 | 35 | ....R..T..... |
| GO:0042393 | histone binding | 3/30 | 12/3454 | 0.0001 | 0.0033 | 0.0028 | 3 | .....D.TE..... |
| GO:0032991 | protein-containing complex | 2/23 | 14/3454 | 0.0036 | 0.0446 | 0.0405 | 2 | .....Y.SD.... |
| GO:0006511 | ubiquitin-dependent protein catabolic process | 2/23 | 15/3454 | 0.0042 | 0.0446 | 0.0405 | 2 | .....Y.SD.... |
| GO:0003724 | RNA helicase activity | 2/23 | 18/3454 | 0.0061 | 0.0446 | 0.0405 | 2 | .....Y.SD.... |
| GO:0003723 | RNA binding | 6/20 | 199/3454 | 0.0006 | 0.0153 | 0.0147 | 6 | .....SYE..... |

### Conservation analysis

Based on unique phosphorylated protein-sites at 5% FLR in *P. falciparum* 3D7, we investigated the existence of their orthologs in other species of *Plasmodium*. Conservation for each site, defined as the proportion of species containing the same amino acid at that position in the multiple sequence alignment, can be found as supplementary files (Supp File 3.). We generated a heatmap (Figure 4) representing the average agreement among sites within each protein. If the site was conserved with respect to the reference 3D7 it was labelled as 1 and 0 otherwise. For each species, a non-conserved site could be the result of having a different amino acid with respect to the reference phosphorylation position, there could be gap in the protein or the orthologue was not found for that species. Then, within each protein and species, the average (proportion of sites conserved) was calculated. In this way, proteins without any sites conserved or when the protein is not found in the species have score=0 and when all phosphosites are conserved, the score = 1. Changes in a site between Ser, Thr or Try was considered not conserved, although scores were also calculated allowing for S <-> T substitutions as “conserved” (not disruptive to S/T phosphorylation), as shown in Supp File 3.

**Figure 4.**
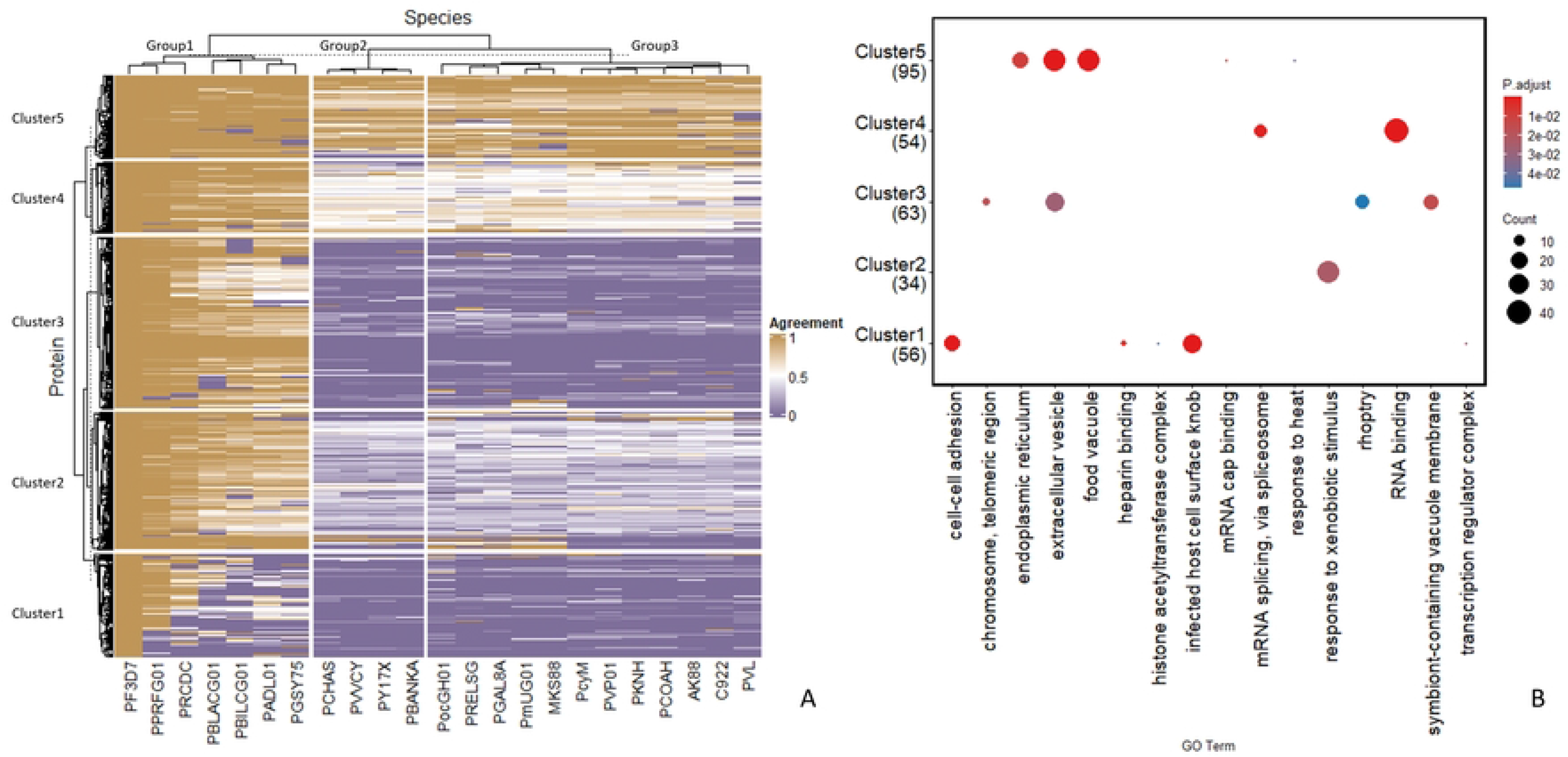
A: Heatmap of the agreement in sequence conservation between 23 species of *Plasmodium*: *P. gorilla* clade G1(PPRFG01), *P. reichenowi* (PRCDC), *P. blacklocki* G01 (PBLACG01), *P. billcollinsi* G01 (PBILCG01), *P. adleri* G01 (PADL01), *P. gaboni* (PGSY75), *P. malariae* (PmUG01), *P. brasilianum* strain Bolivian I (MKS88), *P. ovale* (PocGH01), *P. relictum* (PRELSG), *P. gallinaceum* (PGAL8A), P. chabaudi AS (PCHAS), *P. vinckei vinckei* CY (PVVCY), *P. yoelii* 17X (PY17X), *P. berghei* ANKA (PBANKA), *P. cynomolgi* (PcyM), *P. vivax* (PVP01), *P. knowlesi* (PKNH), *P. coatneyi Hackeri* (PCOAH), *P. fragile* strain nilgiri (AK88), *P. inui* San Antonio 1 (C922), *P. vivax*-like (PVL) and the reference *P. falciparum* (PF3D7). The heatmap displays the mean conservation (agreement) of phosphosites per protein with three clusters for species and five for proteins. B: Pathway enrichment analysis for the genes found in each cluster with significant GO terms in y-axis and clusters in the x-axis; the number of genes where those terms were found are in parenthesis under each cluster label.

Mapping across species returned 26,316 sites belonging to 2,890 proteins, where only 1 mapping occurrence was used per phosphorylated protein-site, i.e. when phosphorylated peptides could match to more than one protein (or very rarely multiple positions within one protein) only one match was used per peptide. Of those 2,890 proteins, only 108 proteins contained all phosphosites that were completely conserved across all species considered. Hence, the heatmap in Figure 4A is formed of the 2,782 proteins for which there were differences between two or more orthologous proteins in the conservation of their identified phosphosites.

The dendrogram suggests three main groups for the *Plasmodium* species on the x-axis, with PPRFG01 (*Plasmodium sp.* gorilla clade 1) closest to the *P. falciparum* reference PF3D7 (Figure 4A). This group (group 1) containing *P. falciparum* has six more species in the cluster, none of which (beyond 3D7) are transmissible to humans. The two other groups are formed of 4 and 12 species. All other species transmissible to humans are included in the largest group (group 3). On the y-axis, five clusters of proteins were formed and subsequently analysed separately with clusterProfiler for enrichment analysis, to determine if there were different biological processes associated with phosphosites of different conservation patterns, which might be indicative of those under different selective pressures. The analysis yielded 16 statistically significant GO terms (Figure 4B).

For Cluster 5 (proteins containing phosphosites mostly conserved across the genus), the most significant GO terms per cluster were: “food vacuole” (GO:0020020), ”extracellular vesicle” (GO:1903561) and “endoplasmic reticulum” (ER) (GO:0005783) – indicating conserved signalling mechanisms related to the infection of host cells. The ER in Apicomplexa is known to be involved in the processing of effector proteins before translocation to host cells [31]. The “food vacuole” is a GO term mostly used in annotation of Apicomplexa proteins related to digestion of the host cell cytoplasm. Cluster 4 contains sites that are highly conserved in Group 1 species, and around 50% conserved in Group 2 and 3 species. Cluster 4 is annotated to be enriched for GO terms related to RNA binding and splicing. This result is somewhat surprising, since one would assume that mechanisms related to transcription would be very highly conserved. Cluster 3 are highly conserved in Group 1 species but have low conservation in groups 2 and 3, with enrichment for GO terms related to symbiont-containing vacuole membrane, rhoptry and extracellular vesicle – we could speculate that this may be indicative of cell signalling related to host-cell invasion, and evolving faster that Cluster 5 proteins, to evade host immune responses. Cluster 2 proteins have highly conserved phosphosites in Group 1 proteins but with average conservation between those in Cluster 2 and 4 for other groups. Only one GO term is enriched for “response to xenobiotic stimulus” (GO:0009410) – a term related to *Plasmodium*’s ability to respond to small molecules from the host (and proteins implicated in drug resistance). Cluster 1 contains proteins with least conserved phosphosites, and had the strongest enrichment (by p-value and count of mapped terms) for “infected host cell surface knob” (GO:0020030) and “cell-cell adhesion” (GO:0098609). Term GO:0020030 is a commonly used gene annotation in *Plasmodium*, related to the protrusions in the membrane of an infected erythrocyte. These proteins are potentially under pressure to evade host immune responses, and thus evolving much faster (and changing their cell signalling mechanisms). Data for this analysis including the clusters can be found in Supp. File 4.

We next implemented a “strict” phosphosite matching process, for the purposes of providing highly likely phosphosites for all *Plasmodium* species aligned, requiring that the phosphosite amino acid matched between *Plasmodium falciparum* 3D7 and the target species (allowing for S <-> T substitutions) and requiring the +1 residue was also matched (as the most important position for phosphorylation motifs). This gives an additional candidate set of 183,134 phosphosites, identified across the *Plasmodium* genus, based on orthologue mapping (Supp. File 5). The multiple sequence alignments for all phosphoproteins are provided in compressed folder (Supp. File 6).

We also investigated conservation based on SNP analysis within the *Plasmodium falciparum* species, using data sets derived from whole genome sequencing of different isolates downloaded from PlasmoDB (Supp. File 7). Out of 26,006 phosphosites, 25,587 did not have any recorded single amino acid variants (SAAVs recorded in PlasmoDB), indicating very high conservation of phosphosites within the species sampled (98.3%). Analysis of the total serines within the proteome found 4,660 SAAVs with major allele frequency (AF) <1 (from 261,791 total serines), i.e. 98.2% conservation. This indicates that that pS is no more likely or less likely to be mutated than other serines. On average, 97.9% threonines in the proteome are conserved (i.e. have no SAAV in this analysis), compared to 98.7% for pT sites. On average, 99.0% of pY (663/670) and 99.0% of all tyrosines do not have a SAAV in this analysis, again indicating no particular selective pressure signal that could be identified. A histogram of the major allele frequencies for phosphosite SAAVs is presented in Figure 5B, confirming that most phosphosites are highly conserved across different isolates, with only a single site (pSer 33 in PF3D7_1366900, a protein of unknown function) having major AF < 0.5. A table of proteins with phosphosites and AF < 0.9 is provided in Supp. File 7, including several zinc finger proteins and two rhoptry proteins.

**Figure 5.**
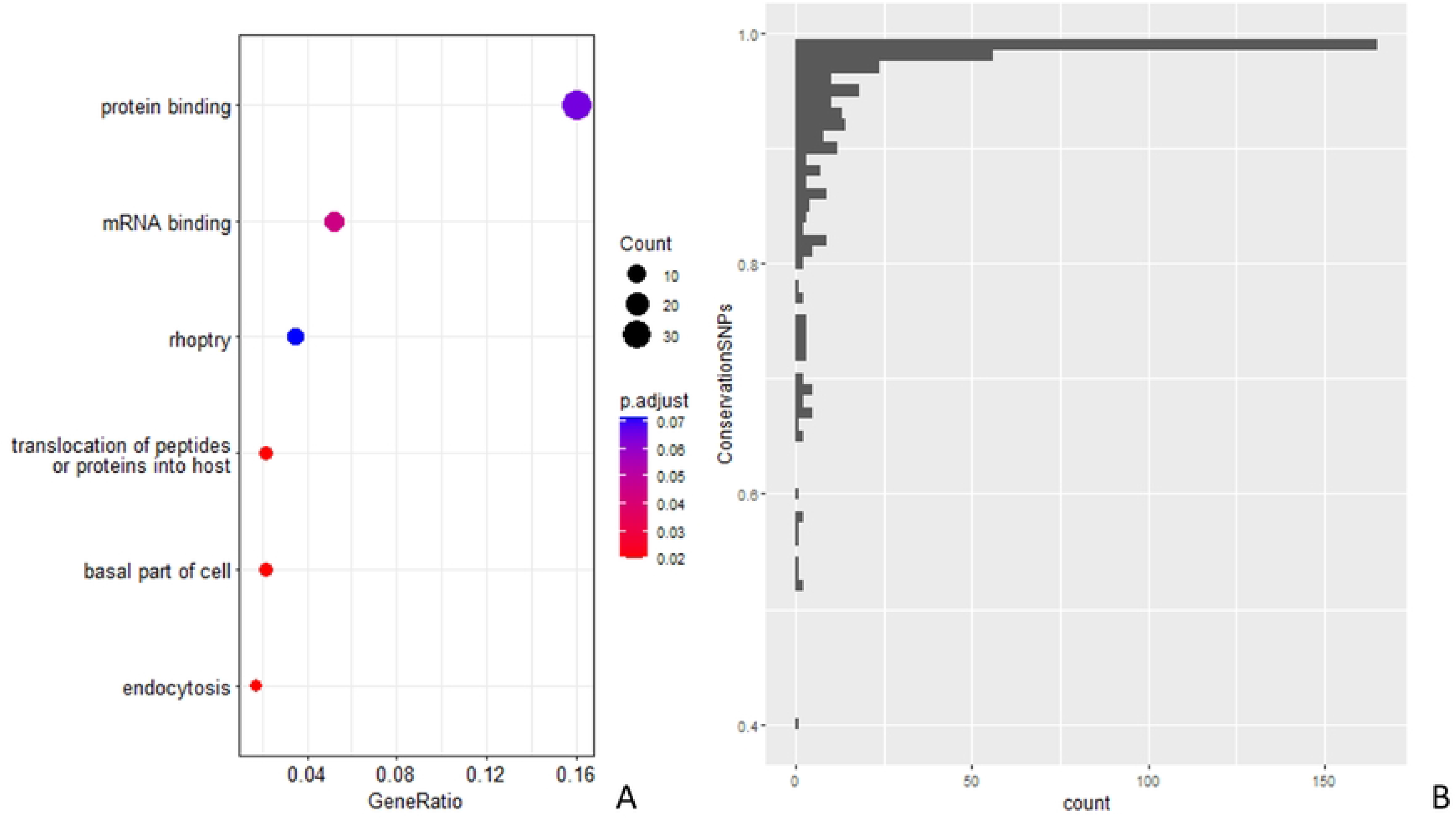
A: Pathway analysis results across all *Plasmodium* species for those genes which were not fully conserved compared to PF3D7, based on SNP analysis. B: Histogram of conservation across *Plasmodium* species based on SNP analysis.

We performed GO term enrichment analysis for the (419) proteins containing not fully conserved phosphosites (Figure 5A). Besides “protein binding” (GO:0005515) or “mRNA binding” (GO:0003723) which appear to be significant in most analyses, the analysis also returned “translocation of peptides or proteins into host” (GO:0042000), “rhoptry” (GO:0020008) and “endocytosis” (GO:0006897), indicative perhaps of proteins under some positive selective.

Other comparative analyses of conservation data have been included as Supp. Figure 2. In Supp Figure 2 (A) a pathway analysis was performed comparing three human transmissible species (PmUG01, PocGH01, PVP01) vs. 17 not transmissible. *P. knowlesi* (PKNH) and *P. vivax*-like (PVL) were excluded from this analysis because of their potential for non-vectorial infection to humans. From phosphoproteins, a subset was selected for pathway analysis as: proteins with fully conserved phosphosites for the human transmissible species, i.e. all phosphosites conserved with respect of the reference PF3D7, and not conserved for non-transmissible species, at least one phosphosite within the protein not conserved with respect to the *P. falciparum* reference PF3D7. Some of the significant terms not previously observed in other analyses were “cytosol”, “structural constituent of ribosome”, “cytosolic small ribosomal subunit”. Supp Figure 2 (B) investigates the enriched pathways for sites within phophoproteins not conserved in *P. gorilla* clade G1 (PPRFG01), compared to PF3D7, as our analysis (Figure 4A) suggested this species to be the closest relative to *P. falciparium*. This additional analysis returned “infected host cell surface knob” (GO:0020030) as the most significant term, which may indicate differences on the cell invasion between the two species of *Plasmodium*.

### Disorder and structural analysis

Next, we explored the structural context of phosphosites in *P. falciparum*, to search for information about the functional importance of the phosphosites. First, we performed an analysis to predict all the structured and disordered regions of *P. falciparum* proteins (using metapredict v2), and mapped phosphosites onto these regions. Metapredict gives a score from 0-1 to indicate the likelihood of each amino acid within a sequence to be in an ordered (score = 0) or within a disordered region (score = 1). It has been reported before that a high proportion of mammalian phosphosites are located on disordered regions, having a role in transition from disorder to order, altering the local or global structure, and potentially changing the interaction potential of the protein [32]. From our mapping of phosphosites to predicted disordered regions, we could observe that phosphosites had a very strong tendency towards disordered regions in *P. falciparum*. This tendency for phosphosites to be found in disordered regions can be observed for all three residues S, T, Y. Figure 6A, displays boxplots with strikingly higher disorder scores for phosphosites (pS, pT, pY) than for other S, T or Y residues in the *P. falciparum* proteome – median disorder scores pS=0.982, pT = 0.965, and pY=0.975 compared to S= 0.769, T= 0.639 and Y= 0.532 median disorder scores of all residues in the proteome. The trend is particularly striking for pY sites, since tyrosines do not have a strong preference to be in disordered regions of proteins, as shown on Figure 6A. Metapredict documentation suggests that a threshold of 0.3 can differentiate ordered from disordered regions. Using such a threshold across the entire proteome of *P. falciparum* would suggest that 66% of all residues fall in disordered regions. Comparing against metapredict disorder scores for the human proteome (Supplementary Figure 3), revealed that only 39% of residues within the human proteome are located in disordered regions. It is possible that *P. falciparum* proteins are generally more disordered than human proteins, or that the tool is less well calibrated for Apicomplexa than for humans. Nevertheless, using the metapredict-recommended threshold of >0.3 for determining disordered regions, in agreement with previous reports [26], revealed that 89% of phosphosites were located within predicted disordered regions (and still 85% if a more conservative score >0.5 was used to determine disorder).

**Figure 6.**
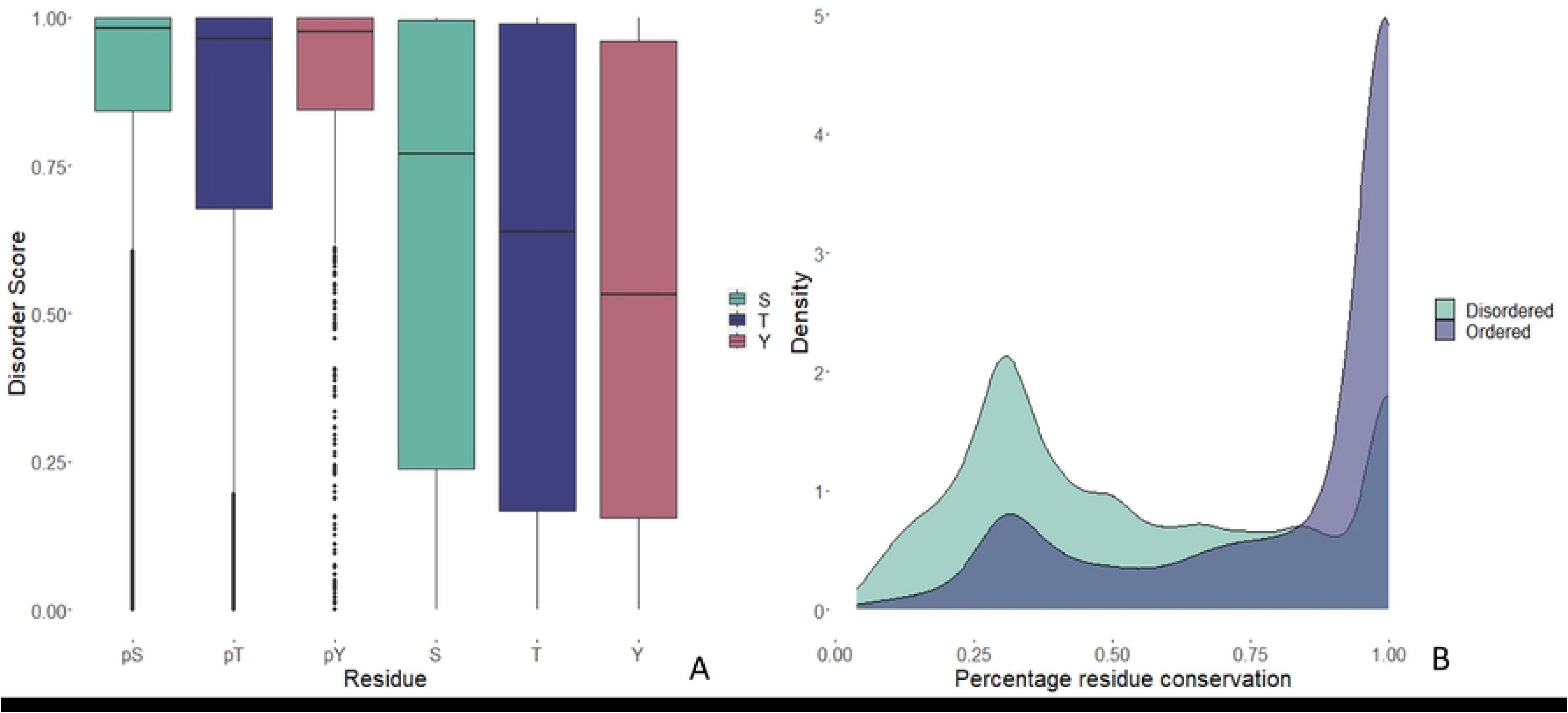
A: Boxplot of the disorder scores (from metapredict) for phosphosites (pS, pT and pY) versus disorder scores for all S, T and Y residues in the *P. falciparum* proteome. B: Density functions for percentage of conservation by residue for ordered and disordered regions (<0.3 metapredict score).

If we restrict the analysis to FLR “Gold” category phosphosites (those observed with high confidence in more than one study), remarkably only 5.8% fall in ordered regions (on only 330 proteins), indicating that it is highly unusual for phosphorylation to occur on well-structured regions of *P. falciparum* proteins. For comparison, 12% of “gold standard” human phosphosites [23] are predicted to fall into ordered regions (metapredict score < 0.3). Investigating the biological functions links to these 330 *Plasmodium falciparum* proteins with phosphosites in their ordered regions, there several GO terms returned from pathway enrichment analysis with clusterProfiler (Supplementary Figure 4), including proteins localising to the cytosol and cytoplasm, and ribosome-related functions.

Next, we wished to explore whether there was any difference in the conservation of phosphosites in disordered vs ordered regions. As shown on Figure 6B, there is a striking difference – phosphosites in ordered regions (2,776), had high conservation overall.

Examining the set of ordered sites, since these are relatively unusual in the *P. falciparum* phosphoproteome, these sites are highly conserved (Figure 6B), compared to disordered phosphosites – 48% of all “ordered” sites have conservation >90% across the genus, compared to only 16% of “disordered” sites.

Exploring the high-quality (Gold) set of sites mapped to ordered regions (330 proteins), 209/330 (63%) have a human ortholog (OrthoMCL DB [33]), indicative of genes highly conserved across all eukaryotes. For the proteins containing “Gold” quality phosphosites in disordered regions (1,724), 718/1,724 (42%) have a human ortholog – indicative of proteins that are less well conserved across eukaryotes. Disorder and site conservation data is available as Supp. File 8.

### Protein structural context

The release of AlphaFold2 (AF2) has had a very significant impact on the ability to understand the three-dimensional (3D) structure of proteins across both model and non-model organisms [34]. 3D structure predictions for *P. falciparum* proteins are available via AlphaFold database, UniProtKB and PlasmoDB (via mapping to UniProt identifiers). We have mapped all the phosphosites onto AF2 structures, also incorporating the conservation scores (across the *Plasmodium* genus), enabling visual exploration of the relationship between order/disorder (which can be clearly visualized) on structures, conservation and positions of phosphosites. In Supp. File 9, we have created a static html page with a table of the phosphoproteins identified, with hyperlinks so that every phosphoprotein’s structure and phosphosites can be visualized via the online iCn3D viewer [35], and links to their corresponding record in UniProtKB (see section below on data access). A caveat is that AF2 models have no awareness of PTM sites, and generally have been trained on few example proteins with PTMs intact. As such, protein structure models demonstrate the position of PTMs as they would appear on an otherwise unmodified structure. Given that phosphosites often change the structure of proteins by introducing more negative charge, it remains an open research question how to re-model AF2 structures to reflect the presence of phosphosites.

An example is presented in Figure 7, for protein phosphatase PPM2 (UniProtKB identifier: Q8IHY0, PlasmoDB identifier: PF3D7_1138500). The image displays all identified phosphosites on the green to red colour scale, indicating fully conserved across the genus = green, to unique to *P. falciparum* = red. In PPM2, it can also be observed that phosphosites that are highly conserved (green) are located in the structured core, and that large, disordered regions are around the outside, containing non-conserved phosphosites in red.

**Figure 7.**
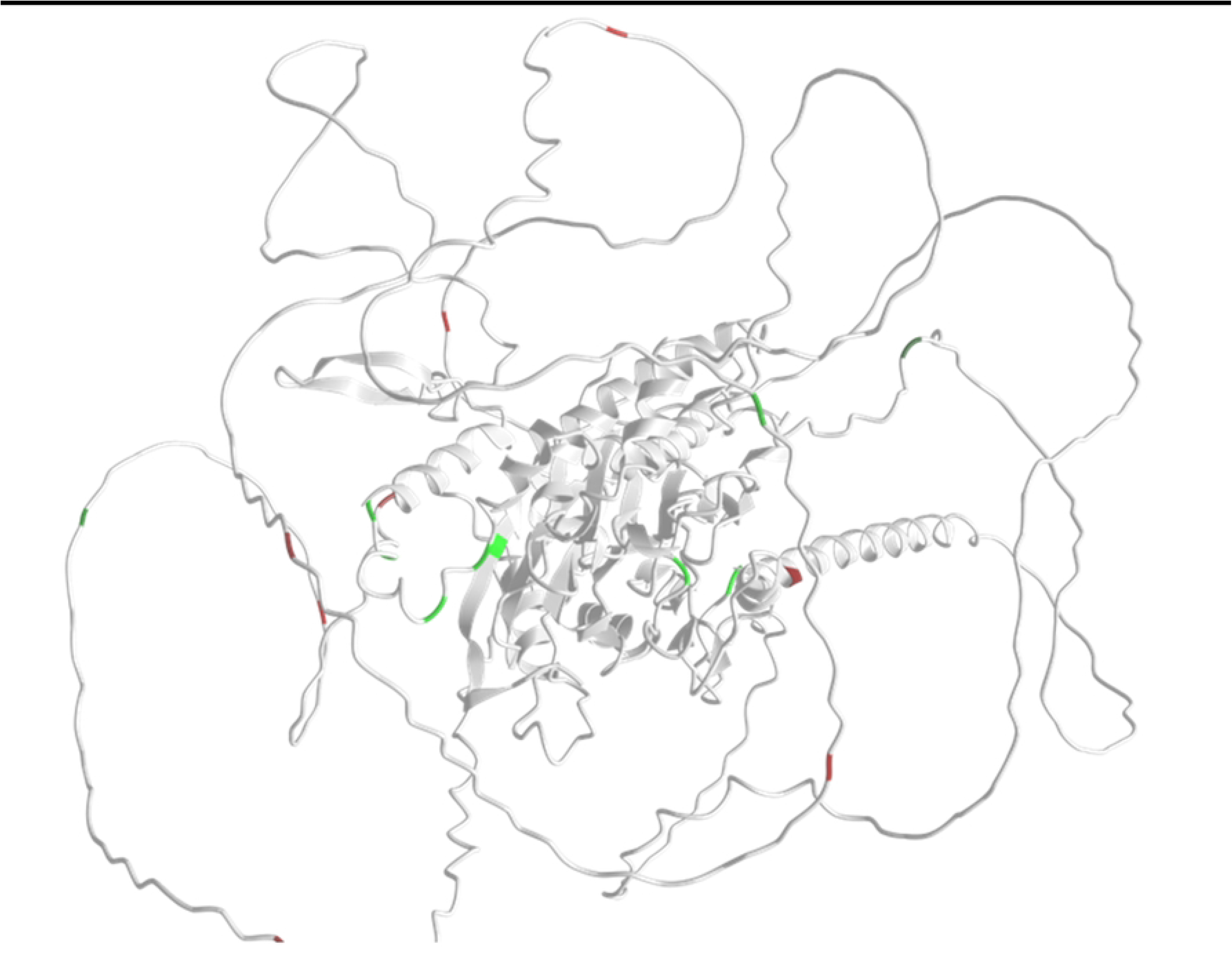
Protein PF3D7_1138500 protein phosphatase PPM2, visualised in iCn3D (AlphaFold structure Q8IHY0), with mapped phosphorylation sites – coloured red-black-green conservation scale (0-9). Phosphosites in the structured core are fully conserved across the *Plasmodium* genus, whereas disordered regions have mostly low conservation.

When exploring functional relationships across orthologues, it might be typical to conclude that highly conserved regions are most significant for function, and in fact most protein domains are captured from conserved regions in multiple sequence alignments across assumed orthologues. However, the data presented here does not support such a conclusion for phosphosites. The fact that the vast majority of phosphosites are present on disordered regions of proteins, which are evolving fastest, and seem to have a role in host cell invasion and potentially evading host cell responses, point to a specialisation in the functional role of phosphorylation. Further experiments to understand the interplay between phosphorylation, protein disorder and the ability to infect hosts are clearly required.

### Open access data availability

The *Plasmodium falciparum* “phosphosite build” is part of a wider project, called PTMeXchange, aimed at high quality re-analysis of MS/MS data sets enriched for particular PTMs, and providing simple public access to the resulting data sets. The build is provided via PRIDE/ProteomeXchange with identifier PXD046874, where researchers can download the phosphosites identified per study in simple tab-separated text files. The data has also been loaded into UniProtKB (Figure 8A), enabling phosphosite evidence to be explored alongside other protein features and AF2 models, with links to the raw evidence. We have also released the phosphosite build within PeptideAtlas (https://peptideatlas.org/builds/pfalciparum/phospho/), enabling more detailed exploration of the evidence within each protein, peptidoform and spectrum for a given site (Figure 8B). Our data are deposited in PRIDE, UniProtKB and PeptideAtlas all provide evidence for each site using Universal Spectrum Identifiers [36], which can be rendered via https://proteomecentral.proteomexchange.org/usi/ (and shown in right panel on Figure 8B). This allows a user to explore the evidence for a given site on a peptide, and test other possible explanations to see if the spectrum could support alternative explanations (peptides or sites within those peptides). Lastly, the phosphosite build is scheduled to appear in PlasmoDB in 2024, as this is the central database for *Plasmodium* researchers.

**Figure 8.**
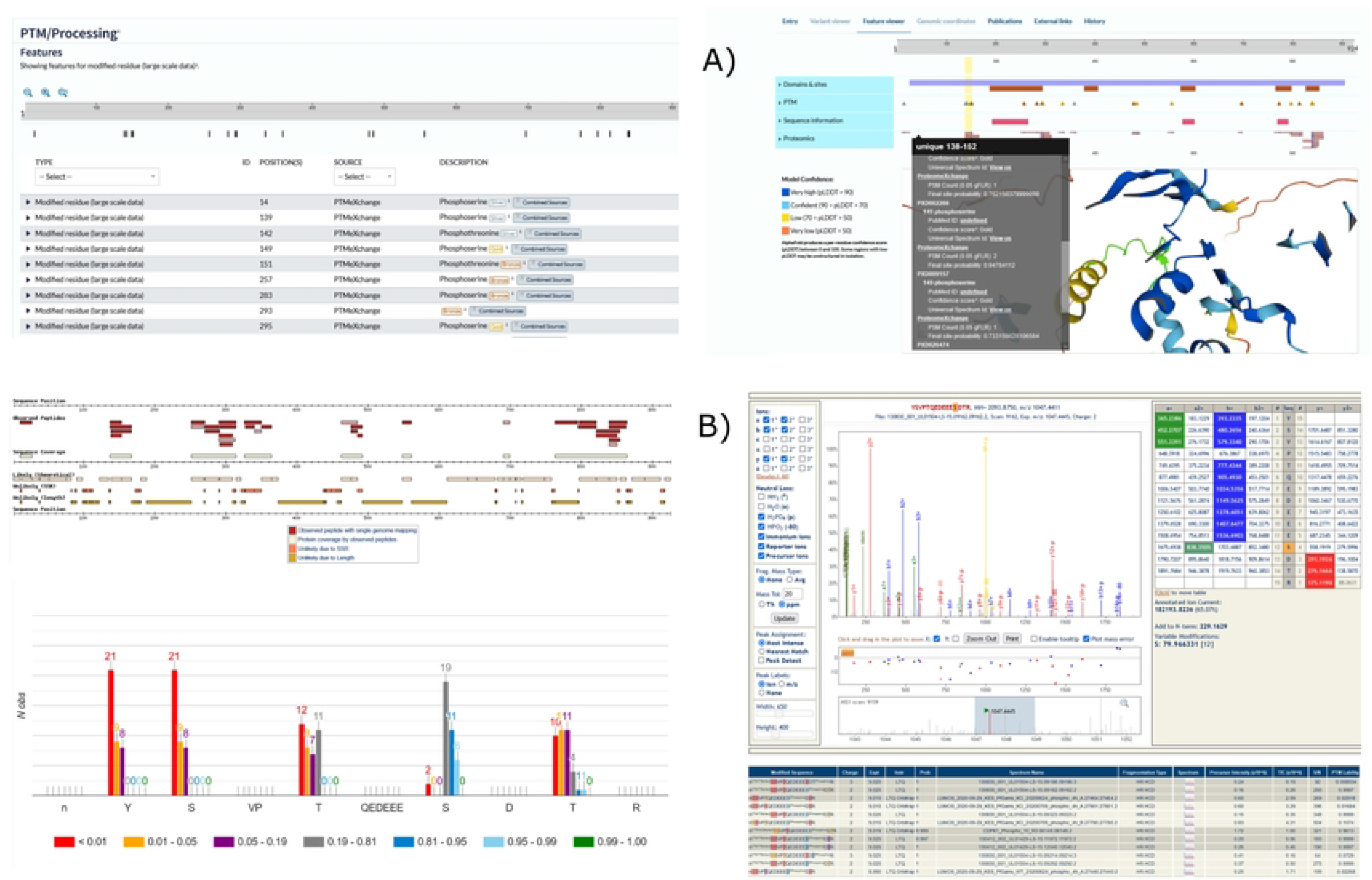
A. Visualisation of phosphosites in UniProtKB for example protein (Q8IHY0). B. Examples of different visualisations at the protein-, peptidoforms-, and spectral-levels for the same protein, with PeptideAtlas identifier PF3D7_1138500.1-p1.

## Discussion

We have reported a comprehensive meta-analysis of *P. falciparum* phosphoproteomics data sets. Data sets were re-analysed using a robust pipeline ensuring objective false localisation rate calculation. Additional Gold, Silver, Bronze labelling was also provided as easy way to display confidence of phosphosites being correctly identified.

Results from our meta-analysis have been deposited into UniProtKB, enabling sites to be used for further research with other bioinformatics tools. The data has also made available in PeptideAtlas and PRIDE, enabling detailed exploration of scores and visualization of source mass spectra, as a full evidence trail.

We have also provided over-represented motifs found in these phosphoproteins as well as conservation data in relation to another 22 species of the genus *Plasmodium*, and from 115 different *Plasmodium falciparum* isolates, with respect to *P. falciparum* strain 3D7. We provided predictive disorder scores for all identified phosphosites and links to protein structures to visualise these proteins. We expect our results to be a useful resource for researchers facilitating identification of areas for future research in developing malaria treatments and vaccines.

## Methods

### Data selection

The ProteomeXchange Consortium [37] was used to identify suitable *P. falciparum* phosphoproteomics datasets, via the PRIDE repository [38] and ProteomeXchange. Overall 11 were deemed suitable for phosphosite (localisation) reanalysis, based on inclusion criteria that data sets were generated by data dependent acquisition (DDA) methods, “high-quality” (i.e. likely to deliver >1,000 phosphosites), raw data were available and readable using open source tools: PXD000070 [39], PXD001684 [4], PXD002266 [40], PXD005207 [41], PXD009157 [42], PXD009465 [43], PXD012143 [44], PXD015093 [45], PXD015833 [30], PXD020381 [46], and PXD026474 [47]. These 11 data sets consist of both labelled (TMT or iTRAQ) and unlabelled MS data sets. Most of the studies focus on the blood stage in *P. falciparum* life cycle, with only one study (PXD026474) including gametocytes. There was not data available for the liver stage of the parasite, possibly due to challenges with the development of liver models. A brief description of the studies’ objectives extracted from the abstracts of their publications, *P. falciparum* stages included in the analysis, enrichment details and MS methods are shown in Table 2 and more details about the studies’ objectives and biological samples in Supplementary Table 1.

### Phosphosite localisation

The search database was created from sequences derived from PlasmoDB 51 release of *Plasmodium falciparum* 3D7 and human. The human sequences were obtained from the Level 1 PeptideAtlas Tiered Human Integrated Search Proteome [48], containing core isoforms from neXtProt [49] (2020 build). A fasta file was created combining these sequences with cRAP contaminant sequences (https://www.thegpm.org/crap/) plus decoy sequences. One decoy sequence was generated each protein and contaminant sequence using the de Bruijn method (with k=2) [50].

Data analysis was performed using the Trans-Proteomic Pipeline (TPP) [19], including Comet [51] search engine for individual datasets, followed by PeptideProphet [52], iProphet [53], and PTMProphet [54]; these were grouped within each study according to the labelling and MS characteristics, as they determine search parameters. The files were searched with several variable modifications depending on the study design, and phosphorylation of ASTY. Alanine was included in searches as a decoy amino acid to enable the estimation of the FLR as explained by Ramsbottom *et al*. [21]. Other variable modifications included in the analyses were: Oxidation in MWH (HYDR), protein N-terminal acetylation or at K (ACET), ammonia loss at peptide n-terminal QC (PYRO), pyro-glu at peptide n-terminal E (DHB), deamidation at NQ (DEAM), formylation of the N-terminus (FORM) and fixed modification Carbamidomethylation (C), as noted in Supplementary Table 1. iTRAQ4plex, TMT6 or TMT10 labelling were included in searches as appropriate. As search parameters, a maximum of 2 missed cleavages specified and maximum number of 5 different modifications per peptide were allowed.

### Post-processing search results

The data files obtained from searching with TPP were processed by custom Python scripts (https://github.com/PGB-LIV/mzidFLR) and analysed following a previously published pipeline [22].

First, at the peptide-spectrum match (PSM) level, the FDR was calculated based on decoy sequences and the PSMs were filtered to 1% FDR. Data from most confident PSMs (1% FDR) were transformed to give a site localisation score for each phosphosite found on each PSM, removing matches to decoy PSMs and contaminant entries. A final site-based PSM score was obtained by multiplying the peptide identification probability by the site localisation probability and adjusted considering the number of occasions each site was observed, phosphorylated and not phosphorylated, in the dataset [22].

Once the final probability at PSM-level was calculated, the data was collapsed to “peptidoform-site” level by taking the maximum score among all matches for each phosphosite on a given peptidoform within each analysis and study. FLR was calculated by ordering all peptidoform-sites by their final score. At this stage, matches to *P. falciparum* and human matches were separated according to their respective protein matches.

A further categorisation at the protein-site level was achieved by using FLR calculations for each study. This Gold-Silver-Bronze (GSB) categorisation allows additional grading for our confidence in the findings among the most confident sites. A simple exclusive criterion was applied: Gold - sites observed at 2 or more independent studies at 1% FLR; Silver - sites observed in only 1 dataset at <1% FLR; Bronze: all other sites at 5% FLR.

Note that within each study, if there was more than 1 peptidoform-site as evidence for a protein site, the one with lowest FLR was used for GSB categorisation.

### Downstream analysis Summarising outcomes

A frequency table was produced for PSMs at 1% FDR level, PSM-sites at 1% FDR level, overall number of phosphosites at PSM-level at 1% and 5% FLR. Human and *P. falciparum* number of sites were separated at peptidoform-site level and protein-site level at 1% and 5% FLR.

GSB phosphosites counts were generated considering that some of the sequences, and therefore sites, could map to different proteins or sites within their respective proteome. The number of phosphosites in sequences mapping to a single site (unique) and to more than one site were also calculated, as well as the number of phosphosites mapping to only either the *P. falciparum* or Human proteome, or belonging to sequences that map to both proteomes.

All analyses and graphical output were obtained using the R programming language (version 4.2.1) or above, via RStudio (2022.02.3 Build 492).

### Motif and pathway enrichment analysis

All *P. falciparum* datasets were investigated using motif and pathway enrichment analysis. For this, we used all STY phosphosites at 5% FLR, combining all 11 studies and removing matches to the decoy amino acid Alanine. 15mer peptides centred on each phosphosite were generated to investigate motifs around these sites. They were compared against 15mer “background” sequences generated using any STY in these datasets, whether they were phosphorylated or not, at its centre (position 0). Statistically significant enriched motifs surrounding phosphosites were identified using the R package *rmotifx* [55]. Results were thresholded via p-value < 1e-9 for pS and p< 1e-6 for pT and pY respectively; and a minimum of 20 sequences per motif.

The R package clusterProfiler [56] was used to carry out a pathway enrichment analysis considering those phosphoproteins containing specific enriched motifs. All other proteins in the search database were used as background for this analysis. A heatmap on the adjusted p-values of a subset of motifs with fold-change enrichment (versus the background) > 4 was produced representing motifs against GO (Gene Ontology) terms.

### Conservation analysis

*P. falciparum* isolate 3D7 identified phosphosites were investigated regarding 22 species of the *Plasmodium* genus. The 22 species of *Plasmodium* were: *P. gorilla clade G1*(PPRFG01), *P. reichenowi* (PRCDC), *P blacklocki* G01 (PBLACG01), *P. billcollinsi G01* (PBILCG01), *P. adleri G01* (PADL01), *P. gaboni* (PGSY75), *P. malariae* (PmUG01), *P. brasilianum strain Bolivian I* (MKS88), *P. ovale* (PocGH01), *P. relictum* (PRELSG), *P. gallinaceum* (PGAL8A), *P. chabaudi AS* (PCHAS), *P. vinckei vinckei CY* (PVVCY), *P. yoelii 17X* (PY17X), *P. berghei ANKA* (PBANKA), *P. cynomolgi* (PcyM), *P. vivax* (PVP01), *P. knowlesi* (PKNH), *P. coatneyi Hackeri* (PCOAH), *P. fragile strain nilgiri* (AK88), *P.inui San Antonio 1* (C922), *P. vivax-like* (PVL). They were compared against *P. falciparum* 3D7 in terms of sequence conservation. Protein sequences were downloaded from PlasmoDB and mapped to the identified phosphosites using the “syntenic ortholog” mappings stored in PlasmoDB, generated by OrthoMCL [33], followed by protein-level multiple sequence alignment with muscle 5.1 running on Linux [57].

Matched residues in orthologs were labelled as 1 while not matching sites, due to having a different amino acid in that position in the ortholog, a gap in the sequence, or simply not having an ortholog for a protein were labelled as 0. For each species and phosphoprotein considered, a mean conservation score was calculated for the proportion of phosphosites within each protein, which were conserved within the orthologue from that particular species. Heatmaps were created based on the mean conservation score. Protein clusters were identified and investigated for enrichment analysis using clusterProfiler.

Further functional enrichment analyses were performed comparing conservation results from human transmissible species *P. malariae*, *P. ovale and P. vivax* to the other 17 animal species (excluding *P. knowlesi* and *P. vivax-like* due to their zoonotic character). The subset of proteins included in the enrichment analysis were those proteins fully conserved for the three human transmissible species and those not fully conserved across all the animal species.

Conservation of phosphosites within *Plasmodium falciparum* was calculated using single nucleotide polymorphism (SNP) data stored in PlasmoDB (public search strategy: https://plasmodb.org/plasmo/app/workspace/strategies/import/b4d2489952494797), using variants from 115 aligned genomes from two unbiased SNP data set https://plasmodb.org/plasmo/app/record/dataset/DS_9a7f849906, and https://plasmodb.org/plasmo/app/record/dataset/DS_d1c8287de9 [58]. SNP sites were downloaded and non-synonymous variants, altering the amino acid sequence, were matched to the phosphosite positions in proteins (using Python 3.9 code). Site conservation was estimated using the major allele frequency, calculated by PlasmoDB.

We next explored the average disorder of amino acid positions within proteins using metapredict v2 [59] running on Linux, comparing and correlating disorder scores with site classifications. To visualise examples of phosphosites with different conservation values, we used iCn3D [35]. The list of *P. falciparum* 3D7 kinases was generated by searching for proteins annotated with Interpro or Pfam “protein kinase” domains in PlasmoDB: https://plasmodb.org/plasmo/app/workspace/strategies/import/6a11331d6eea4d20).

Phosphosite data for *Plasmodium falciparum* 3D7 was loaded into UniProtKB, by taking all mapping all peptides carrying GSB sites to proteins in UniProtKB *P. falciparum* 3D7 proteome (https://www.UniProt.org/proteomes/UP000001450), assuming tryptic cleavage.

## Acknowledgements

We are grateful to all the researchers that generated MS data and generously shared via PRIDE/ProteomeXchange allowing this analysis.

## Author Contributions

O.M.C. performed database searching, data analysis and manuscript writing. K.A.R. performed search database generation, assisted with data processing. A.P. supported MS data curation. Y.P.R. supported PRIDE data upload and curation. J.F., and M.M. assisted with data loading into UniProtKB and E.B-B visualisation in UniProtKB. E.W.D and Z.S. assisted with MS data searching and creation of the PeptideAtlas build. J.A.V. assisted with PRIDE data loading and project supervision. A.R.J. coordinated the research, assisted with data analysis and writing the manuscript.

## Supplemental Information

Supp File 1. GSB_withMotifs.csv : Phosphosites at 5% FLR classified as Gold, Silver and Bronze and statistically significant motifs for those sites.

Supp File 2. EnrichmentResult_Allmotifs_05.xlsx : Enrichment analysis results for all statistically significant motifs.

Supp File 3. Conservation.csv : Conservation scores across species with respect to the reference 3D7.

Supp File 4. ConservationSpeciesWithClusters.csv : Proportion of sites conserved within each protein and species with respect to 3D7 and its clusters.

Supp File 5. mapped_plasmodium_sites.tsv : Phosphosites putatively identified in other *Plasmodium* species based on orthologue mapping.

Supp File 6. plasmodium_alignments.zip : Multiple sequence aligments of orthologous phosphoproteins within the *Plasmodium* genus.

Supp File 7. SNP_data.xlsx : Conservation analysis based on single amino acid variants.

Supp File 8. Disorder.csv : Disorder scores from metapredict for Gold, Silver and Bronze phosphosites. Supp File 9. Proteins_3D : Hyperlinks for visualising identified phosphoproteins in iCn3D viewer.

Supplementary Information.docx : Suplementary tables and figures.

